# The Human Cerebellum: A Digital Anatomical Atlas at the Level of Individual Folia

**DOI:** 10.1101/2024.08.27.610006

**Authors:** John G. Samuelsson, Jeremy D. Schmahmann, Martin Sereno, Bruce Rosen, Matti S. Hämäläinen

## Abstract

Scientific interest in the cerebellum has surged in the last few decades with an emerging consensus on a multifaceted functionality and intricate, but not yet fully understood, functional topography over the cerebellar cortex. To further refine this structure-function relationship and quantify its inter-subject variability, a high-resolution digital anatomical atlas is fundamental. Using a combination of manual labeling and image processing, we turned a recently published reconstruction of the human cerebellum, the first such reconstruction fine enough to resolve the individual folia, into a digital atlas with both surface and volumetric representations. Its unprecedented granularity (0.16 mm) and detailed expert labeling make the atlas usable as an anatomical ground truth, enabling new ways of analyzing and visualizing cerebellar data through its digital format.

## Introduction

The last few decades have seen a paradigm shift in cerebellar neuroscience and neurology. While the cerebellum has historically been viewed mainly as a motor control center, neuroimaging and clinical case studies have established the cerebellum as a structure containing critical nodes in the distributed neural networks that govern sensorimotor, autonomic, cognitive, and emotional processing with an intricate topography (Albus, 1971; Andersen et al., 1992; Andreasen and Pierson, 2008; Herculano-Houzel, 2009; Marr, 1969; Middleton and Strick, 2001; Raymond and Medina, 2018; Schmahmann, 1991; Schmahmann et al., 2019; Schmahmann and Sherman, 1998; Sereno et al., 2020; Shambes et al., 1978; Stoodley and Schmahmann, 2009).

Along with our advancing understanding of the wide range of roles that the cerebellum plays in health and disease, having access to adequate tools with which to examine cerebellar neurophysiology has become increasingly important. One such tool is a digital atlas that can be used to specify and associate anatomical regions of the cerebellum with certainty, allowing for data integration across subjects. Substantial progress has been made over the past 25 years in the development of cerebellar atlases designed for this purpose. For almost 40 years the definitive atlas of the human cerebellum was that of Angevine et al. (1961). The development of the Talairach atlas (Talairach and Tournoux, 1988) that made it possible to map the human brain in coordinate space using magnetic resonance imaging provided detail of the cerebral hemispheres, but no information about the cerebellum.

The branch point in atlasing of cerebellum for the contemporary era commenced with the publication by Schmahmann et al. (1999) followed up by the more comprehensive MRI Atlas of the Human Cerebellum (Schmahmann et al., 2000). This atlas placed the cerebellum in the Montreal Neurological Institute (MNI) coordinate space, identified the cerebellar lobes, lobules and fissures, and clarified and simplified the nomenclature that had vexed the field for 200 years. The Schmahmann Atlas made it possible to view activations on task-based MRI in the cerebellum with anatomical detail not previously available. It also enabled development of the SUIT atlas (Diedrichsen, 2006) and subsequent, progressively more sophisticated elaborations of probabilistic and multimodal atlases as well as automated volumetric segmentation algorithms based on advances in computer vision and machine learning, which are now widely used in task-based, resting state functional connectivity, tractography and morphometric MRI studies of the human cerebellum (Buckner et al., 2011; Carass et al., 2018; Diedrichsen et al., 2009; Han et al., 2020; Hernandez-Castillo, 2023; Jenkinson et al., 2012; Lyu et al., 2024; Makris et al., 2003; Nettekoven et al., 2024; Ren et al., 2019; Romero et al., 2017).

As useful and transformative as these atlases have been, there is no available digital cerebellar atlas with resolution fine enough to capture folia-level structures that encompasses both volumetric and cortical representations. The availability of such an atlas would be extremely valuable for a wide range of purposes. Some applications include using the cortical surface to constrain the inverse problem in M/EEG source estimation or fMRI analysis. The surface can also be used to visualize neuroimaging data in inflated and flat representations, and a user can interact with these manifolds using a rendering software. The volumetric representation could also be used together with a subject-specific segmentation to register the atlas to a subject, and then applying the registration transformation to the surface, thus achieving a more reliable surface reconstruction than what is possible using only a cerebellar mask.

To address this unmet need in cerebellar atlasing, we set out to develop a digital atlas of the cerebellum to the level of the cerebellar lobule and individual folia, and that incorporates both volumetric and cortical representations. The regions of the atlas are listed in **Table 1**. The cortical atlas representation with 32 regions was further subdivided into 815 smaller patches. The deep cerebellar nuclei are not represented in the present atlas.

**Table 1.**
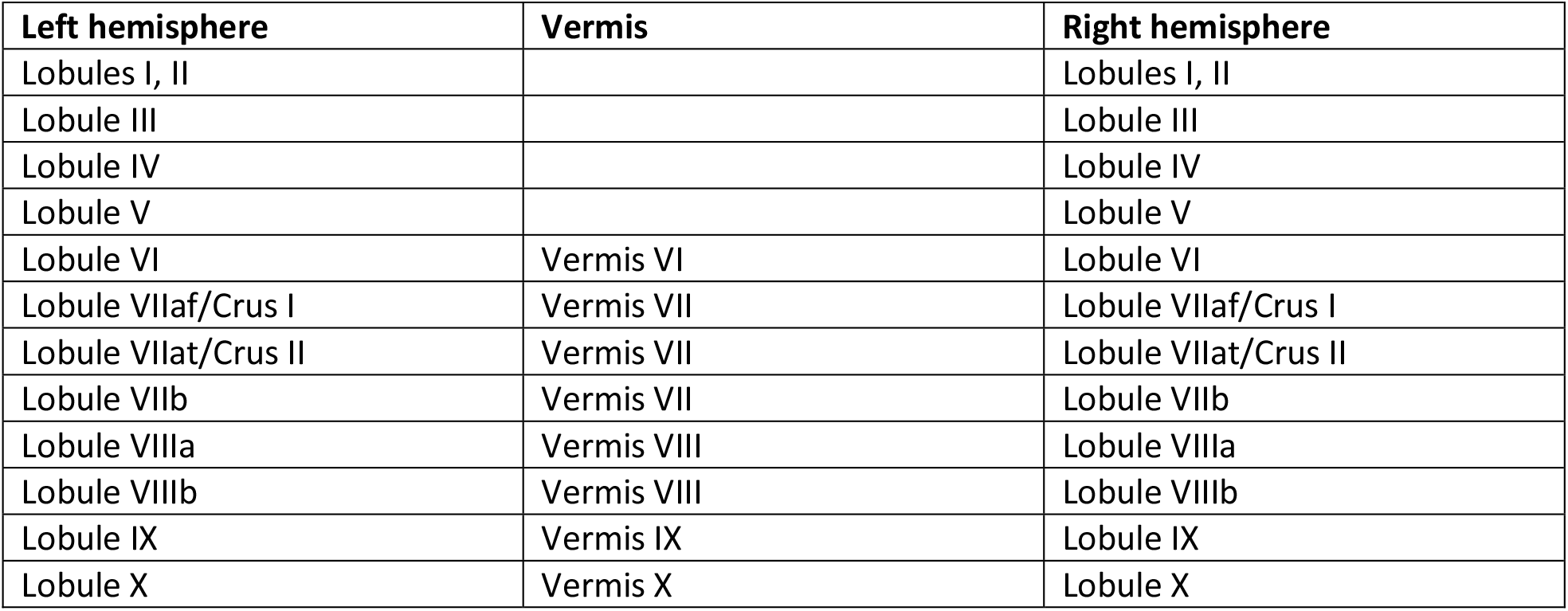
Regions of the cerebellar atlas.

## Methods

The atlas was created by a combination of manual annotation and image processing of the first surface reconstruction of the human cerebellum at the level of individual folia, presented in Sereno et al. (2020). The spatial resolution of the reconstruction, and thus also the derived atlas presented here, is about 0.16 mm in edge length between neighboring vertices in the surface representation. The volumetric representation of the atlas was calculated from the surface representation at an approximate virtual isotropic voxel resolution of about 0.4 mm. An overview of the process for creating the cerebellar atlas is outlined in **Figure 1**, and the detailed account follows.

**Figure 1.**
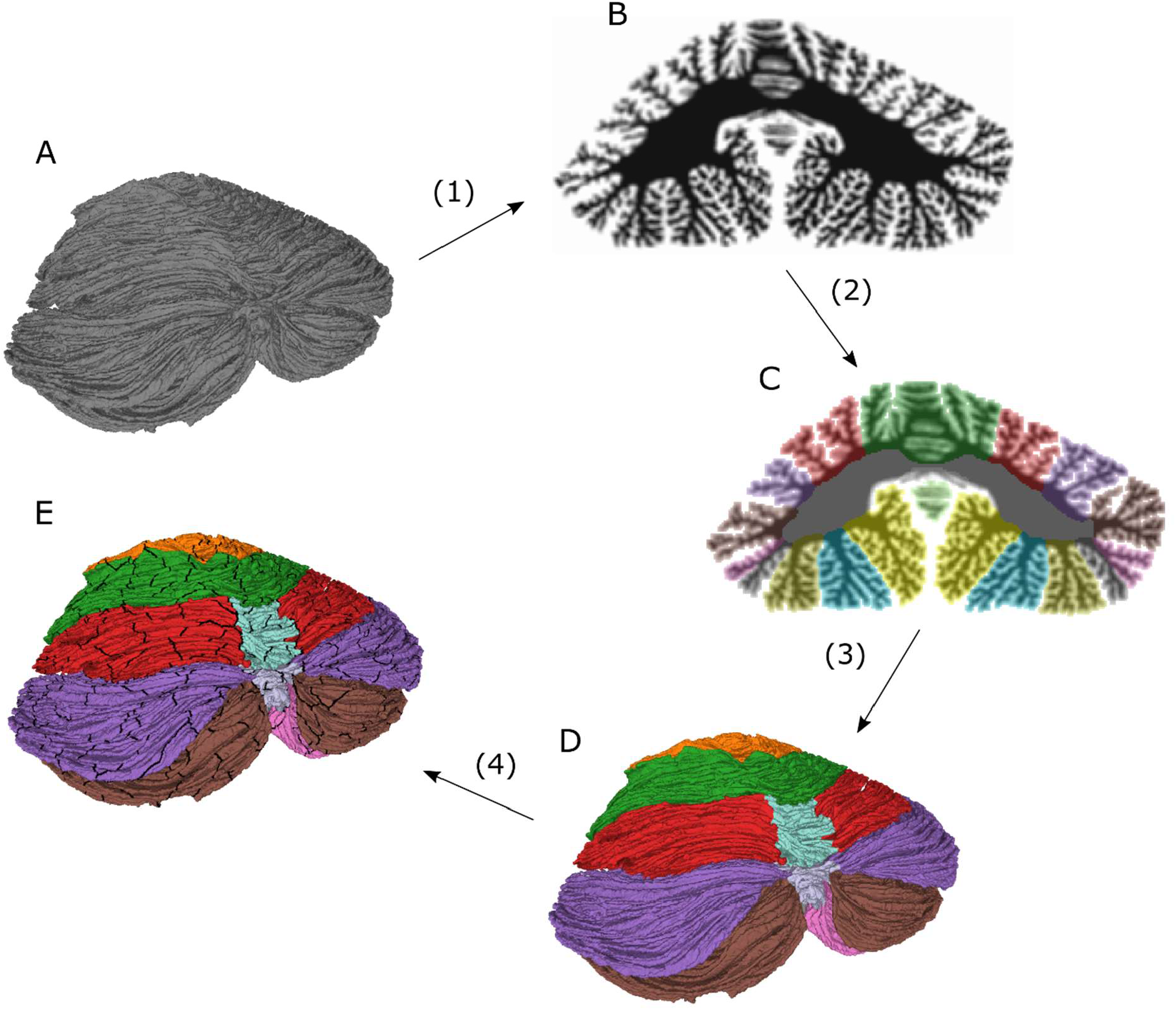
Conceptual schematic outlining the methods for creating the cerebellar atlas and surface parcellation. 1) Starting from the original surface (A), a virtual volume (B) is created by computing solid angles with the surface at points in a regular volumetric grid. 2) The volume is manually annotated using Freeview software and processed with correction and smoothing filters (C). 3) The volumetric atlas (C) is turned into a surface atlas (D) by assigning each vertex in the surface tessellation to the voxel label in which the vertex resides. The surface atlas is processed with a surface smoothing filter and the consistency between the surface and the volumetric atlas is ensured by labeling each voxel as belonging to the mode of the vertex labels in each voxel. 4) The surface atlas is further subdivided into 815 non-overlapping patches, resulting in a surface parcellation (E).

### 1) Creating a virtual MRI volume

Starting from the surface tessellation containing about 4.6 million vertices and about 9.2 **million** triangular faces in Sereno et al. (2020), the surface was down-sampled to about 1.1 million vertices to reduce computation time while working with the atlas (the final atlas was at the original resolution, containing about 4.6 million vertices). A virtual volume was created by computing the solid angles with the surface at a set of observation points (apexes) in a regularly, isotropically spaced rectangular grid. The apexes were about 0.4 mm apart. The value of the solid angle was either 4π, 2π, or 0, corresponding to an apex location inside, on, or outside the surface, respectively. From the solid angle data, artificial voxels were generated by first dividing the solid angles by 4π, so that each apex attained a value of 1, 0.5, or 0. A set of virtual voxels was then created by centering a voxel at each apex. A virtual contrast was computed for each voxel as the sum of the solid angle of the apex at the center of the voxel and the 26 neighboring apexes in the rectangular volumetric grid, so that the virtual contrast would be roughly proportional to the portion of the voxel that lay inside the surface (1 = voxel completely inside surface, 0 = completely outside). This contrast was then multiplied by 2, making it an integer between 0 and 54, so that the data could be saved as an unsigned 8-bit integer and thus reducing storage space.

### 2) Manual annotation and image processing of volume

The virtual volume was manually annotated in the sagittal and coronal planes using Freeview software (Fischl, 2012), one cross-section at a time. In cases where the delineation between lobules was unclear, the cross-section was moved to a point where the delineation could be asserted with higher confidence, and the sulci that made up the division was then followed through the cross-sections, in line with our current understanding of cerebellar neuroanatomy. The labeled volume was saved, and loaded into Python 3.7 using Nibabel version 3.1.0 (Brett et al., 2020) and all annotations that had been made by mistake at voxels outside the cerebellar cortical surface, i.e., where the virtual voxel contrast was computed to be 0, were discarded. The voxels that were not completely outside the volume (with contrast > 0) that had been missed in the annotation was iteratively assigned the mode of its neighboring voxels until all voxels that were not completely outside the surface had been labeled. A volumetric smoothing filter was then applied in three iterations where each voxel in the volume was assigned the mode label of its neighbors to avoid individual voxels that had been labeled by mistake, and to smooth the borders between neighboring anatomical regions. The resulting atlas was carefully manually reviewed to ensure consistency with the Schmahmann atlas.

### 3) Turning volumetric annotation into surface

A surface atlas was created by assigning to each vertex the voxel label in which the vertex resided. Any vertex that was inside a non-labeled voxel (with contrast = 0) was assigned the label of the closest vertex that had been labeled. The surface atlas was then smoothed to avoid small patches of non-uniform labeling where the cortical surfaces of different topographical regions were close. The smoothing was done in three iterations by assigning the mode label of all vertices connected to each vertex by three edges or less. Finally, to make sure that the volumetric and surface atlases were consistent, all voxels that contained any vertices were assigned the mode label of all vertices contained within each voxel.

### 4) Parcellation of the surface atlas

The surface atlas, containing 30 distinct anatomical regions, was further parcellated into 815 patches of approximately equal size (mean=1.9, std=0.55, min=0.83, max=3.5 [cm^2^]). This was done by first defining the number of patches in each region to 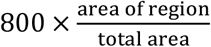 for an initial parcellation of 800 patches. The number of patches was chosen so that the area of the patches would be approximately equal to the patches in the Laussanne parcellation at the finest scale (Cammoun et al., 2012); the Laussanne parcellation contains 1000 patches and the cerebellar cortex is roughly 80% of the area of the cerebral cortex (Sereno et al., 2020).

A number of vertices equaling the number of minor patches defined as explained above was then randomly chosen in each major patch. These vertex pointers were then moved away from one another in an iterative fashion. This was done by moving each pointer from its vertex to the neighboring vertex that was furthest away in terms of geodesic distance and that belonged to the same anatomical region, thus spreading the pointers over the cortical manifold. The geodesic distance was approximated as the Euclidean distance in a spherical surface representation of the surface mesh. This was done for each pointer in 100 iterations.

The patches were then created by recursively expanding each patch from the vertices associated with each pointer by including each neighbor of each edge vertex into the patches. This expansion continued until all vertices had been assigned to a patch. To make the patches more circular and avoid narrow strips, the surface division was smoothed using the same technique described above for the surface atlas; each vertex was labeled as the mode label of all vertices connected to that vertex by three edges or less in three iterations. All patch areas were then calculated and the patches with a surface area larger than 3.1 cm^2^ were split using the FreeSurfer split function while all patches smaller than 0.83 cm^2^ were merged with the neighboring patch with which it shared its longest border. This was done in an iterative fashion. This procedure did not converge for all patches and a final iteration was made where all large patches were split once and all small ones subsequently merged, resulting in a final total of 815 patches.

## Results

The atlas has both a volumetric and a cortical surface representation. The surface representation is further parcellated into 815 non-overlapping small patches, as described in the methods section. The surface atlas is shown in **Figure 2** from different angles along with a sagittal cross-section around the midline and a coronal cross-section where the cuts have been filled with the corresponding volumetric atlas data. The bottom row shows enlarged images of different sections to elucidate the resolution of the surface representation. In Sereno et al. (2020), the surface manifold was divided into larger anatomical regions that were flattened. Inflated and spherical representations of the surface were also included. Because there is a one-to-one mapping between the vertices in these different representations, the surface atlas was transferred onto the inflated and flat representations. These are shown in **Figure 3** along with the parcellation where the boundaries between the small patches are marked in black.

**Figure 2.**
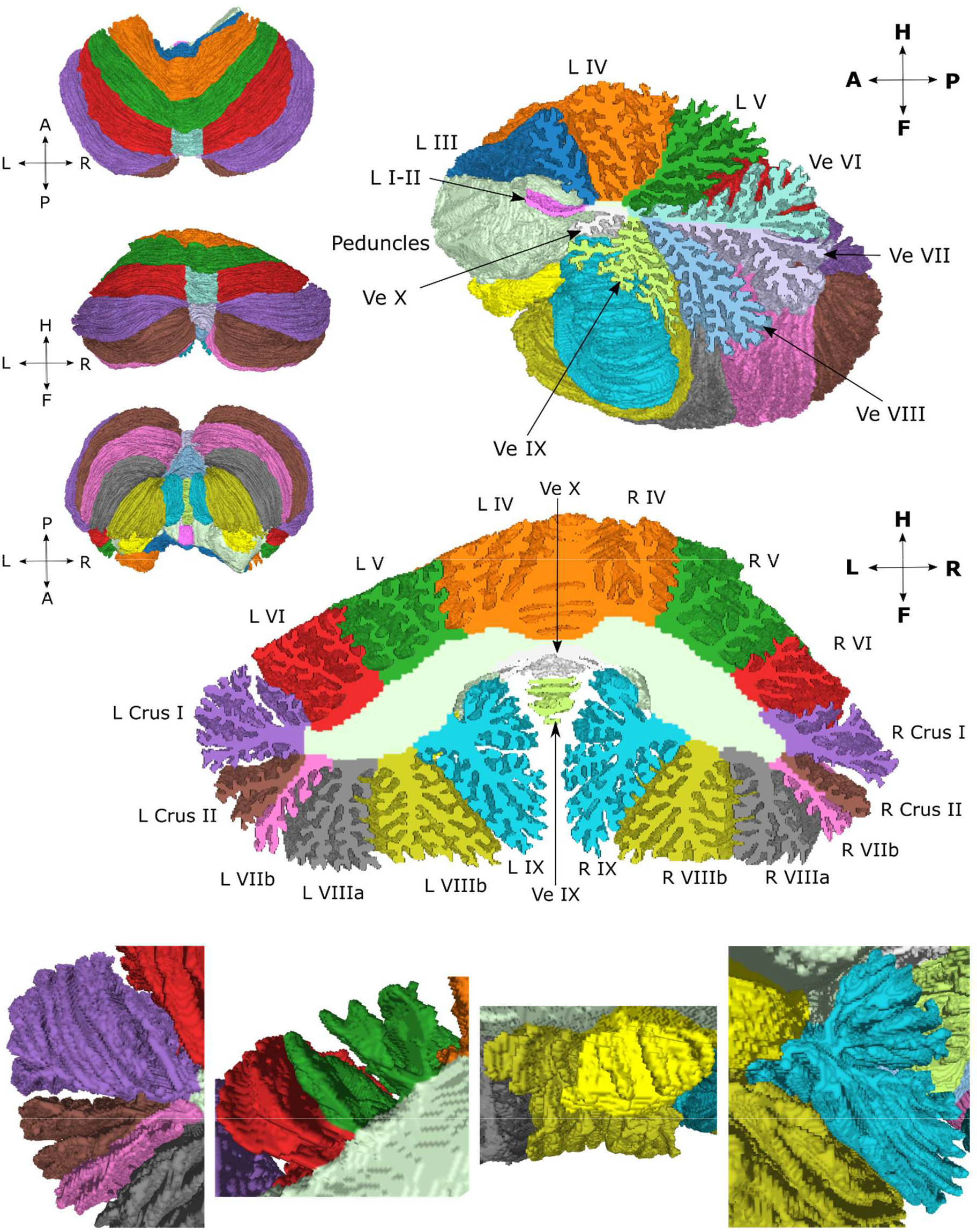
Surface atlas from different views along with a sagittal and coronal cross-section that have been filled with the volumetric atlas. The bottom row shows enlarged images of the surface atlas from the anterior view of left crus I, II and lobule VIIb (left image), lobules V and VI (center-left image), lobules X and VIIIb (center-right image), and lobule IX (right image).

**Figure 3.**
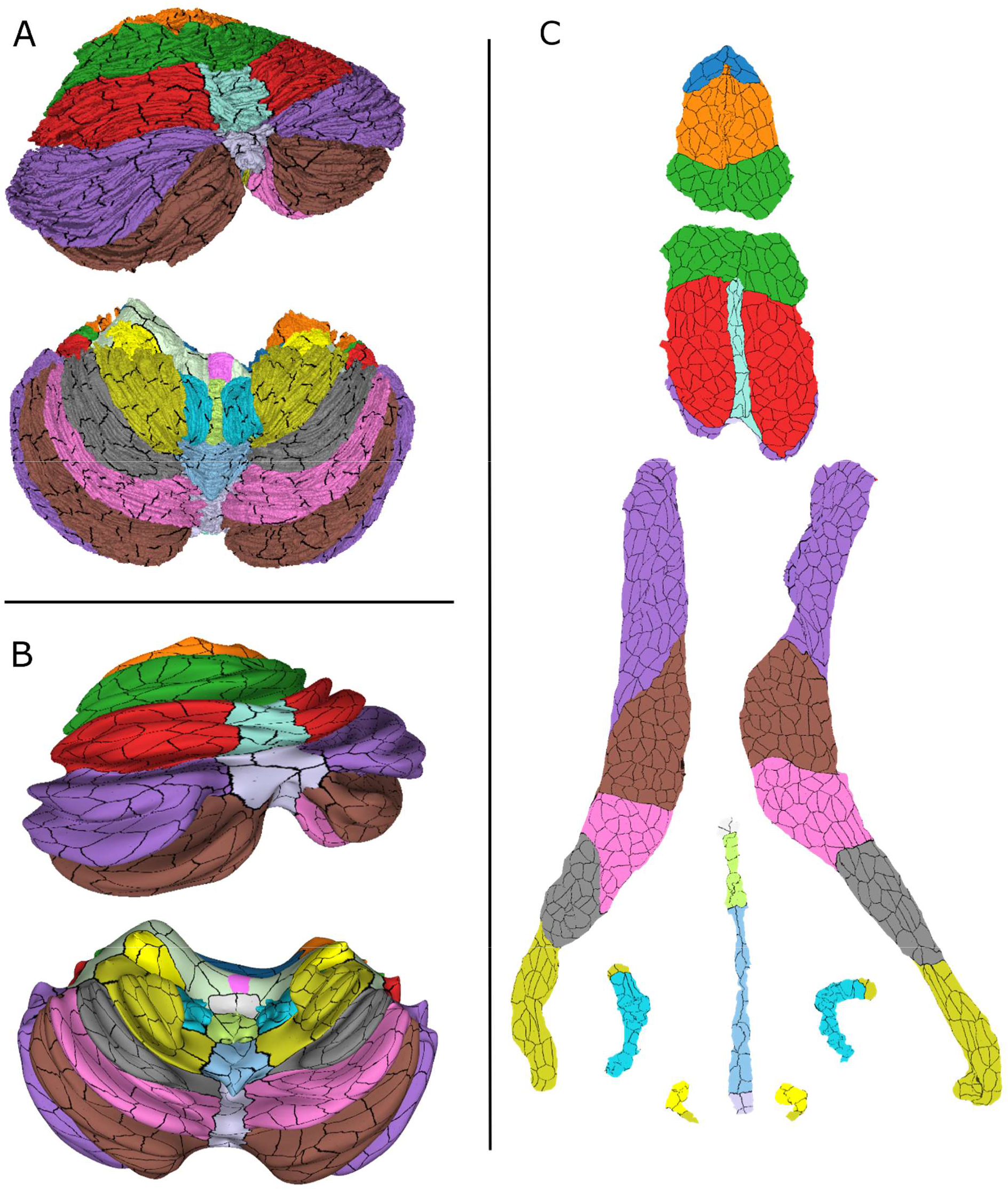
Surface atlas with parcellation borders marked as black in different representations: normal (A), inflated (B) and flat (C).

Our surface atlas is delineated by the cerebellar cortex and makes a closed surface by including the cut surfaces of the superior, middle and inferior cerebellar peduncles and a small stretch of their exposed stalks in the fourth ventricle, annotated here as white matter tracts. The nomenclature is derived from the Schmahmann Atlas, the regions of which are listed in **Table 1**.

## Discussion

Human cerebellar atlases have been pivotal in the transformative studies that have led to deeper understanding of the cerebellum and its multiplicity of functions in the nervous system (Schmahmann et al., 2019). The contemporary topographical divisions of the cerebellar cortex that were employed in this study (Table 1) date back to the early descriptions of the cerebellum (Bolk, 1906) and were reaffirmed with extensive comparative morphological studies (Larsell, 1958; Larsell and Jansen, 1972) and detailed anatomical and physiological investigations in experimental models (Chambers and Sprague, 1955; Jansen and Brodal, 1954; Oscarsson, 1965; Voogd, 2003). As the range and complexity of cerebellar functional incorporation into the distributed neural circuits subserving neurological function and the precision of the topographic arrangement of cerebellar connections and functional topography have become increasingly apparent, the need for a digital cerebellar atlas that enables both surface and volumetric representations with resolution high enough to resolve the anatomy of the cerebellar cortex at the level of the individual folia has become more pressing.

Here we have aimed to address this gap by utilizing the recent high-resolution surface reconstruction of the cerebellar cortex presented in Sereno et al. (2020). This reconstruction was transformed into a virtual MRI volume which was manually annotated, corrected using the image processing scripts described in the Methods section Steps 1 and 2 (Figure 1), and reviewed for anatomical accuracy in comparison with the Schmahmann Atlas. The volumetric annotation was then transferred to a surface annotation in accordance with Step 3 (Figure 1). Finally, each surface annotation was divided into 815 non-overlapping small patches, resulting in a cerebellar surface parcellation in accordance with Step 4 (Figure 1). The resulting atlas and surface parcellation are presented in Figures 2 and 3.

There are many uses of a digital atlas. First, it provides a means for the user to interact with the atlas in a 3D rendering software. Using our atlas, it is possible to rotate, zoom, slide or choose to see specific regions of the atlas, providing an intuitive way of understanding cerebellar anatomy at a high granularity level. There are other digital volumetric atlases such as SUIT (Diedrichsen, 2006) that also allow for such interactions using rendering software, but the highly detailed surface representation available in this atlas in addition to the volumetric representation enable new possibilities such as registration of the volumetric atlas to the segmentation of the cerebellum of a single individual and subsequent application of the registration transformation to the vertices of the surface, and thus morphing the surface atlas to the subject space, yielding an approximate but labeled and high-resolution reconstruction of the cerebellar cortex for that subject. This approach has already proved successful in early demonstrations in our lab (Alho et al., 2023; Samuelsson et al., 2020a). Such surface reconstructions are essential in studies that use a cortical manifold, e.g., in M/EEG source estimation or fMRI analysis (Samuelsson et al., 2020b). Using such an approach, our atlas could enable more spatially specific analysis, making it possible, for example, to compute surface-constrained M/EEG signal power in a specific lobule or patch, or explore the topography of cerebro-cerebellar connectivity with greater precision than possible with current imaging software and atlases. Furthermore, the digital atlas enables an intuitive visualization of topographic data of the human cerebellar cortex *in vivo* since it has inflated, flat and spherical representations.

In annotating the atlas, we encountered and revisited two major anatomical issues discussed explicitly in Schmahmann et al. (1999) and Schmahmann et al. (2000). First, the authors discussed what they called the problem of the vermis, noting that there is indeed no true ‘‘vermis’’ in the anterior lobe. They noted that the application of this term to the paramedian sectors of the anterior lobe term is an extension of the Latin term (meaning worm) used by Malacarne (1776) to denote the structure visible in the posterior and inferior aspect of the cerebellum. While the vermis is morphologically distinct only in lobules VII-X, its presence in lobule VI and the anterior lobe is nonetheless by now well established in the literature. In this atlas, we chose a pragmatic approach of letting the vermis extend from lobule VII throughout lobule VI and end in the primary fissure, keeping the width of the vermis approximately constant throughout vermis VI in the lack of an anatomical landmark to delineate its lateral boundaries. We also created a second atlas where vermis lobules VI and VII are allocated to the hemispheres, so that the vermis encompasses only the vermis of lobules VIII, IX and X. This atlas is not shown but available for download.

The second issue that we revisited was the definition of cerebellar lobules according to the arrangement of the cerebellar fissures. The lobules of the cerebellum are delineated by deep fissures between separate bundles of folia that cross the midline from one hemisphere to the other. This approach builds on the seminal observations by Bolk (1906) and Larsell (1958) and was crystallized in Schmahmann et al. (1999) and Schmahmann et al. (2000). In the present study we observed that one folium, separating crus II from lobule VIIb, violated this paradigm by crossing lobule boundaries around the vermis. It was already noted in Sereno et al. (2020) that the axes of individual folia are often tilted with respect to the overall axis of a lobule; Sereno et al. (2020), Figure 4, shows folia near the base of several lobules progressing toward lobule crests as one moves medially. The alternative of strictly adhering to the paradigm of letting only fissures delineate lobule boundaries, and therefore annotating a folium as belonging only to lobule VIIb or crus II without a change of label in the vermis, yielded substantial asymmetry in the size of lobule VIIb and crus II between the left and right hemispheres (**Appendix Figure 1**). This recapitulated the same issue addressed and reviewed in Schmahmann et al. (1999) and Schmahmann et al. (2000), which was found to reflect a challenge dating back over a century. The ansoparamedian fissure delineates the boundary between crus II and lobule VIIb, but this distinction has been controversial. It was first proposed by the Basle Nomina Anatomica (1895), although the influential work of Bolk (1906) regarded crus II and lobule VIIb as “lobulus ansiformis, crus II”. The distinction of crus II from lobule VIIb resurfaced in Kappers et al. (1936) and then in Jansen and Brodal (1958) and the exhaustive work of Larsell (Larsell, 1958; Larsell and Jansen, 1972). The availability of cross-referencing three-dimensional MRI in humans *in vivo* in Schmahmann et al. (1999) and Schmahmann et al. (2000) made it possible to define with certainty that the ansoparamedian fissure differentiates crus II from lobule VIIb. As depicted and discussed in Schmahmann et al. (1999) and Schmahmann et al. (2000), crus II and lobule VIIb can indeed exhibit asymmetric sizes across the two cerebellar hemispheres, although this has generally not been included in available human MRI cerebellar atlases.

The ready synopsis of these anatomical and nomenclature pitfalls is that whereas crus I is always separated from crus II by the horizontal fissure, and lobule VIIb is always separated from lobule VIIIa of the pyramidal lobule by the prepyramidal fissure, the ansoparamedian fissure that separates crus II from lobule VIIb can be difficult to ascertain with certainty, as we noted again in this brain. An “ansoparamedian lobule” that includes both crus II and lobule VIIb was discussed by Larsell and von Berthelsdorf (1941), but we do not recommend this designation, preferring the agnostic approach of acknowledging that if there is indeed asymmetry across the hemispheres of crus II and lobule VIIb, this invites future exploration of potential functional and clinical relevance of the observation.

### Limitations

The presented atlas focuses on the cerebellar cortex. The cerebellar nuclei and the fiber tracts in the cerebellar white matter, the *corpus medullare*, have not been annotated. While these could potentially be added in future versions on the atlas, the power of the present analysis lies in the unparalleled detail of the morphometry of the reconstructed cerebellar cortex.

As in Schmahmann et al. (1999) and Schmahmann et al. (2000), we studied a single brain. This approach inherently includes idiosyncrasies of any particular individual’s cerebellum morphometry, but it holds the power of defining discrete anatomical features that could be functionally and clinically relevant but potentially overlooked when using averaging and smoothing algorithms necessary for the study of large numbers of brains. The goal of this study was to create a ground truth atlas at a previously unattained resolution that can be used as a template and reference in future studies.

The atlas is based on the surface reconstruction that was presented in Sereno et al. (2020), in which the geometry of the human cerebellar cortex was reconstructed to the level of all individual folia for the first time. Thus, the limitations described in Sereno et al. (2020) also apply to the present atlas. This includes potential drawbacks of reconstructing a specimen *ex vivo*, e.g., tissue swelling and the potential morphological differences introduced when the tissue is removed from the hydrostatic pressure of the cerebrospinal fluid in the intact skull followed by fixation-induced shrinkage.

The finer cortical parcellation presented in this work that subdivides the 30 regions of the atlas into 815 smaller patches was not based on anatomical or functional criteria as was done, e.g., in Makris et al. (2003) and Makris et al. (2005), but rather on a semi-stochastic algorithm to create a set of non-overlapping patches of roughly equal size. The finer parcellation should therefore be viewed mainly as a tool to more precisely describe locations of functional or anatomical data gathered by other means (e.g., MEG/EEG source estimation) than would be possible with the original 30 region atlas.

## Conclusion

We have developed a new anatomical digital atlas with surface and volumetric representations based on the first reconstruction of the human cerebellum to the level of individual folia. As with the advanced atlases of the cerebral cortex, the atlas has unfolded and flattened planar representations. Through its unprecedented granularity, detailed manual labeling and digital format, this atlas may be used as an expert-annotated anatomical ground truth, enabling new ways of analyzing and presenting anatomical and physiological data in the human cerebellum.

## Data availability statement

All atlas and parcellation data are available for download at https://osf.io/98p3a/?view_only=933654b10152444992b9e7d8ff9f1112 and the original full resolution volume and surface data are available at https://pages.ucsd.edu/~msereno/cereb and the image processing scripts are available from the corresponding author upon reasonable request.

## Acknowledgements

This research was funded by the NIH Neuroimaging Training Program (NTP) grant 5T32EB001680, NIH/NIBIB P41 Center for Mesoscale Mapping (1P41EB030006), NIH/NINDS grant 5R01NS104585, the NIH BRAIN Initiative F-32 Fellowship grant 1F32MH127789, NIH Grant R01 MH081990 and UK Royal Society Wolfson Fellowship (M.I.S.). Supported in part by the National Ataxia Foundation, the Raynor Cerebellum Project and the MINDlink Foundation (JDS). The content is solely the responsibility of the authors and does not necessarily represent the official views of the National Institutes of Health. The authors declare that there is no conflict of interest regarding the publication of this article.

## Appendix

**Appendix Figure 1.**
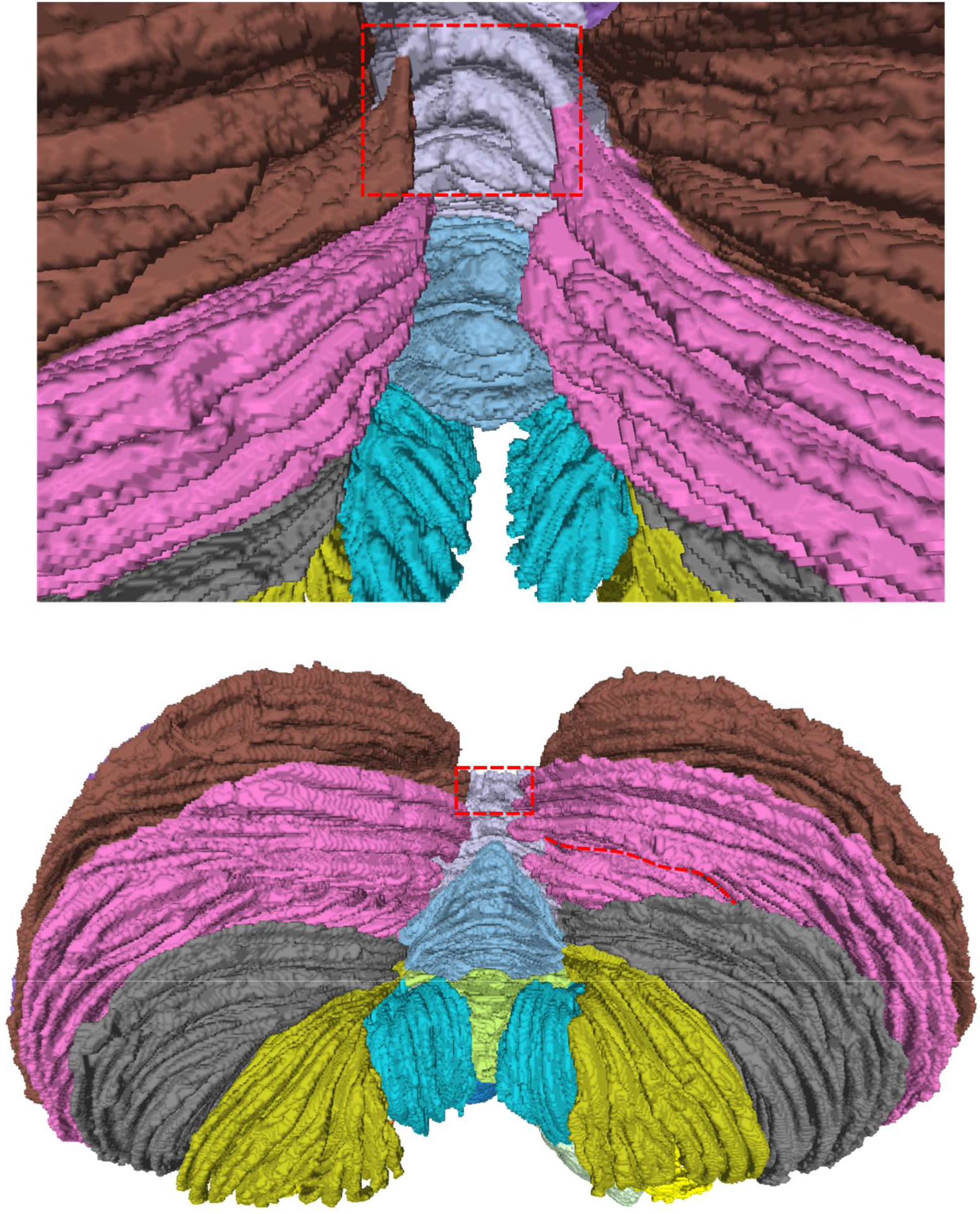
Posterior-inferior view of the surface atlas. The part where the folium of crus II turns into lobule VIIb has been highlighted. In the lower image, the right hemisphere extension of the sulcus that separates lobule crus II from lobule VIIb in the left hemisphere has been highlighted in red. This highlights the substantial lateral asymmetry that would arise by keeping strictly to the paradigm of letting sulci demarcate all lobule boundaries.

